# Tissue atlas of *Cryptosporidium parvum* infection reveals contrasts between the natural neonatal calf model and laboratory mouse models

**DOI:** 10.1101/2025.05.05.652177

**Authors:** Peyton Goddard, Thomas Tzelos, Beatrice L Colon, Paul M Bartley, Lee Robinson, Leandro Lemgruber, Michele Tinti, Grant MJ Hall, Sarah Stevens, Louise Gibbard, Robert Bernard, George Tytler, David Smith, Frank Katzer, Mattie Christine Pawlowic

## Abstract

*Cryptosporidium* is an apicomplexan parasite that causes diarrhoeal disease. The species *C. parvum* is zoonotic and causes significant morbidity and mortality for both humans and farm animals; most commonly, calves and lambs. A One Health approach that integrates human, animal and environmental health perspectives is required to tackle this disease. Current treatments are limited and ineffective, meaning there is an urgent need to develop new anti-cryptosporidials both for human and animal health. The neonatal calf model is a natural model of infection employed as a tool for drug discovery or generating parasite material. However, the model is seldom utilised to investigate host-parasite interaction. Fundamental information about this model, including the location of the parasite in the gut, is lacking. It is also unclear how the more commonly utilised immunocompromised mouse models of cryptosporidiosis compare to the neonatal calf model. To address this, we established an acute, moderate experimental *C. parvum* infection in neonatal calves. Using transgenic parasites, we created a tissue atlas of infection for neonatal calf gut and immunocompromised mouse models and mapped and quantified infection to draw robust comparisons between models. *Cryptosporidium* infection was observed at high levels throughout the neonatal calf gastrointestinal tract and was not limited to the ileal-cecal junction, as previously suggested. This infection pattern is most similar to the acute cryptosporidiosis mouse model, interferon-gamma knockout mice (IFNγKO). Infection with transgenic parasites allowed us to perform *in vivo* and *ex vivo* tissue imaging of the chronic cryptosporidiosis mouse model, NOD SCID Gamma KO (NSG) mice. In contrast, in NSG mice infection is low in the small intestines and highest in the caecum and colon. Understanding the true distribution of infection in the gastrointestinal tract of these three key animal models provides new perspectives on how to interpret and design drug efficacy studies and provides new insight into host-pathogen interaction.

## 1. Introduction

*Cryptosporidium parvum* is a zoonotic parasite and the causative agent of cryptosporidiosis, an enteric disease affecting both humans and farm animals. For both, the young and immunocompromised are most at risk of severe disease. The parasite is faecal-orally transmitted; zoonotic infection occurs through direct contact between people and animals, or ingestion of food or water contaminated with oocysts. *Cryptosporidium* parasites infect the epithelial cells of the gastrointestinal (GI) tract, causing diarrhoeal symptoms that results in mortality and morbidity, including malnourishment and growth stunting. The Global Enteric Multicenter Study (GEMS) identified *Cryptosporidium* as second only to Rotavirus as the leading cause of moderate to severe diarrhoeal disease in children under two^1^. In human disease, cryptosporidiosis is estimated to cause 7.8 million disability-adjusted life years lost (DALYs) globally^2^. This is echoed in cattle, with the Veterinary Investigation Diagnosis Analysis (VIDA) annual report identifying cryptosporidiosis as the most commonly diagnosed cause of enteritis in neonatal calves^3^. Symptoms of bovine cryptosporidiosis are similar to human disease, which include: profuse, watery diarrhoea (cattle scours), dehydration, and stunted growth. *Cryptosporidium parvum* infects multiple species of livestock, but cattle disease is of particular interest due to the economic impact. In the UK there is an estimated average loss of ∼£128 per severely infected beef calf based on weight loss alone (when weight is compared to that of uninfected animals)^4^. Farm animals are also a frequent documented source of *C. parvum* human infections^5^. Cattle shed large quantities of oocysts into the environment, posing risks of transmission to other livestock and humans, especially in the case of contaminated water supplies^6^. Therefore, a One Health approach that integrates research from a human, animal, and environmental health perspective is critical to tackle this devastating disease.

Treatment and prevention options for both human and animal cryptosporidiosis are lacking. The single drug approved for human use, nitazoxanide, is not effective in those most at-risk of disease^7^, and no vaccine is available. The two licensed drugs for cattle, halofuginone lactate and paromomycin^8^, are recommended for treatment only during early onset of disease, and neither drug has proven effective in eradicating disease. For cattle, a new maternal vaccine was recently licensed^9^. Whilst a recent evaluation of the efficacy of the vaccine indicates a reduction in diarrhoeal episodes^10^, the vaccine only mildly reduces oocyst shedding and is only effective in severe disease cases. This has appreciable limitations for reducing agricultural outbreaks and limiting environmental contamination. Concerningly, drug resistance is also already an emerging threat, evidenced by spontaneous resistance observed when experimentally infected cattle were treated with anti-cryptosporidial compounds^11,12^. Future drug development will need to thoughtfully consider the impact of drug resistance in both human and cattle disease.

Despite the fact that calves are an important host species for *C. parvum,* they are most often employed as a research tool, rather than an infection model to study disease biology. Cattle are almost exclusively used to propagate wild type *C. parvum* for downstream use in water and food quality assurance or in laboratory research. Calf infection models are also utilised to assess the efficacy of pre-clinical compounds in development for human disease^13^. Aside from this, experimental studies of bovine cryptosporidiosis are numerous, yet often focus on the agricultural or environmental impact, rather than host-pathogen interaction. Unlike the immunocompromised mouse models most commonly used in parasitology research, cattle are the natural host. In addition, unlike mice, cattle experience symptoms of diarrhoea. Studies of host-pathogen interaction in cattle stand to illuminate the role of the host immune system in controlling infection and the mechanisms of diarrhoea which are important in disease spreading.

The most fundamental knowledge gap in bovine cryptosporidiosis is the location of infection in the calf GI tract. The prevailing hypothesis is that infection in cattle is restricted to the ileo-caecal junction, a small stretch of tissue that joins the small and large intestine, directly before the caecum^14^. However, this observation lacks experimental evidence. In prior studies, *Cryptosporidium* parasites have been observed in other regions of the small intestine by microscopy^15,16^ but infection has never been robustly mapped or quantified. Therefore, the true breadth and burden of infection are unknown. Pinpointing the site of infection is critical for future studies of host-pathogen biology.

We present the first to our knowledge reported passage of transgenic reporter parasites in the neonatal calf. By exploiting a transgenic reporter strain, we conducted a first-of-its-kind study to map and quantify infection throughout the calf GI tract. We compare the calf infection model to immunocompromised mouse models. We describe a tissue atlas of infection in calves and mice and provide the biological foundation and statistical power to inform future studies.

## 2. Materials and Methods

### 2.1 Animal Ethics

All mouse studies were conducted at the University of Dundee. All experiments were carried out under the authority of licences granted by the UK Home Office under the Animals (Scientific Procedures) Act 1986. Prior to submission to the Home Office, applications for project licenses were approved (WEC2018_03 and WEC2023_02) by the University Welfare and Ethical Use of Animals Committee, acting in its capacity as an Animal Welfare and Ethical Review Body as required under the Act.

All neonatal calf studies were conducted at Moredun Research Institute. Experimental designs were approved prior to the commencement of each study by the Moredun Research Institute Animal Welfare Ethical Review Board (AWERB). Experiment numbers: Passage 1 E25/22, Passage 2 E18/23, Passage 3 E11/24.

### 2.2 Molecular cloning and CRISPR design

Oligonucleotides were purchased from Sigma Aldrich/Merck (see **Supplemental Table 1**). Plasmids were constructed using HiFi cloning (New England Biolabs, NEB) and DNA sequence confirmed by Sanger sequencing (Genewiz).

A reporter cassette was designed for parasites to express fluorescent proteins during different parasite life cycle stages. The *CpEnolase* (cgd1_1960) promoter drives expression of mNeonGreen, referred to hereafter as mNeon, throughout all life cycle stages^17^. A 2A peptide also drives expression of a NanoLuciferase-Neomycin resistance fusion (NLuc-Neo^R^). Expression of mScarlet-I-3xMyc reporter is driven by the *Cryptosporidium* Oocyst Wall Protein 1 (*CpCOWP1*) promoter^18^; female parasites express this red fluorescent protein. We cloned the promoter of a candidate male gene (an AP2 transcription factor; cgd6_2670) and used it to drive expression of mTagBFP2. Unfortunately, this element of the reporter did not work, and we could not detect expression of blue fluorescent protein by any parasites. Henceforth, we refer to this strain, Δ*tk::mScarlet-mNeon-Neo^R^,* as FS3N (<u>F</u>emale <u>S</u>carlet, m<u>N</u>eon, <u>N</u>anoLuciferase, <u>N</u>eo^R^; **Supplemental Figure 1**).

Thymidine kinase (cgd5_4440; *CpTK)* was targeted for replacement with this reporter cassette as previously described, using 50 bp of homology from directly upstream and downstream of the open reading frame^19^.

### 2.3 Generation of *C. parvum* reporter strain

Wild type *C. parvum* (Iowa II strain) oocysts were purchased from Bunchgrass Farms (Idaho, USA). Oocysts were excysted, transfected, and used to infect Interferon gamma knockout (IFNγKO; female, 6 weeks old). Mice were gavaged with a saturated sodium bicarbonate solution 5 minutes prior to infection with transfected sporozoites. Paromomycin sulphate was administered in the drinking water (16 g/L) throughout infection to select for transgenic parasites. Faecal samples were collected and pooled by cage, unless reported otherwise.

DNA was extracted from wild type oocysts or mouse faecal samples using the Quick-DNA Faecal/Soil Microbe Miniprep Kit (Zymo Research); at least five rounds of freeze-thaw were performed prior to sample lysis to maximise extraction of DNA from oocysts. PCR was performed with primers designed to detect integration of the reporter cassette at the *CpTK* locus (**Supplemental Table 1**).

### 2.4 Mouse Infections

IFNγKO mice were purchased from Jackson Laboratory (B6.129S7-IfngtmlTS/J, JAX 002287) and bred in-house at the University of Dundee. Non-obese diabetic/severe combined immunodeficiency disease (NOD SCID gamma, NSG) mice (NOD.Cg-*Prkdc^scid^Il2rg^tm1Wjl^/SzJ*) were purchased from Charles River Laboratories (#614NSG) or Jackson Laboratories (#005557). Paromomycin sulphate for use in mouse studies was purchased from Carbosynth (#AP31110).

#### 2.4.1 Propagation of Reporter Strains

Mice were infected by oral gavage and paromomycin was administered in the drinking water (16 g/L) for all passages. Mice were infected with FS3N strain (described in **2.2**, **Supplemental Figure 2** and **Supplemental Table 2**) or *Δtk::*mNeon-Neo^R^ reporter strain (previously described^20^). Faecal samples were collected and shedding levels measured by NLuc assay as indicated (**Supplemental Figure 2B**). Where larger quantities of oocysts were required, intestinal tissue were collected after euthanasia, the lumen flushed with an excess of PBS, and the content collected and added to faecal samples for oocyst purification. This refinement often significantly increases the oocyst yield and reduces the number of animals required to obtain sufficient parasites.

#### 2.4.2 Mouse Tissue Atlas

Two cages of IFNγKO mice and two cages of NSG mice were used (4 mice per cage, mice 8-9 - week-old males). All animals were administered paromomycin in drinking water throughout the study. One cage of each mouse strain was infected with FS3N oocysts; the second cage of each mouse strain remained uninfected. Infection was monitored by faecal NLuc (described **2.7**). When NLucs reached at least 5×10^4^ RLUs/mg (**Supplemental Table 3**), mice were euthanised for tissue collection.

Five locations along the GI tract (duodenum, jejunum, ileum, caecum, colon) were selected for sampling. Approximately 2 cm lengths of tissue were removed from each location and stored at −20 ^°^C prior to further processing. Sampling order was consistent between animals. Infection of tissue quantified as described in **2.7**.

#### 2.4.3 In vivo imaging of NSG mice

NSG mice (males aged 8-10 weeks) were infected with the *Δtk::*mNeon-Neo^R^ reporter strain^20^. Pooled and individual faecal output was collected as indicated and infection levels monitored via faecal NLuc assay.

At various timepoints post-infection, infection levels were also measured using *in vivo* imaging with an IVIS Lumina LT Series III system (Revvity). 10 ml/kg of D-luciferin (prepared at 15mg/ml; Perkin Elmer, #122799) was administered intraperitoneal (IP) injection. Mice were imaged inside an isolation chamber (XIC-3, Perkin Elmer) inside the IVIS. The induction chamber and isolation chamber were charged with isoflurane simultaneously with vaporised 5% v/v isoflurane (and 2.5 - 3 L/min oxygen). Mice were anaesthetised in the induction chamber and once fully anaesthetised, transferred to the isolation chamber. Isoflurane was maintained at 2-3% (with 1 L/min oxygen). 15 minutes post-luciferin injection images were collected (fluorescent image exposure time set to auto; binning set to small; F/stop at 1; FOV at D for >2 mice, FOV at C for 1 mouse). Images were normalised using the Living Image software (Revvity).

#### 2.4.4 Ex vivo imaging

When infection levels reached approximately 1×10^6^ to 3×10^6^ Relative Luminescence units (RLUs) /mg, as determined by individual faecal NLucs, mice were euthanised. The GI tract was dissected and flushed with PBS. Clean GI tract was placed in a square Petri dish. 1 ml D-luciferin (15 mg/ml) was diluted in 25 ml Dulbecco’s Phosphate Buffered Saline (DPBS); this solution was used to fill and cover the GI tract. Tissue was imaged in the IVIS 15 minutes after administering D-luciferin (fluorescent image exposure time set to auto; binning set to small; F/stop at 1; FOV at D). Image normalisation was carried out in the Living Image software (Revvity), using the Image Adjust tool palette to amend the aggregate colour scale minimum to the minimum colour scale value across all images (p/sec/cm^2^/sr) and the maximum to the maximum colour scale value across all images (p/sec/cm^2^/sr). Total flux (p/sec) was calculated using the Living Image software (Revvity), by selecting a controlled ROI per day per mouse. The mean ± SD of all mice (*n =* 4) were plotted.

### 2.5 Neonatal Calf Infections

All calf infections were carried out at Moredun Research Institute. Neonatal calves were sourced by Moredun Research Institute from local farms that had been free from recent *Cryptosporidium* outbreaks. Calves were enrolled 1-2 days post-birth. Calves were not screened for exposure to or infection with *Cryptosporidium* prior to enrolment in the study.

All animals were given 2.5 ml Parofor Crypto (paromomycin sulphate; Huvepharma) per 10 kg body weight (35 mg paromomycin sulphate per kg body weight), delivered in milk once daily, throughout the study. Treatment with Parofor Crypto selects for growth of the transgenic strain in experimentally infected animals. Treatment also reduces the risk of unintended infection with wild type parasites naturally acquired before calves are transported to Moredun Research Institute and enrolled in the study.

Male calves were infected and fitted with a harness and bag system to collect faecal material^21^. This setup is suitable for use with male calves, since the female anatomy causes urine contamination of faecal samples. This necessitated that all animals for this work be male. Infection was monitored throughout via microscopy as previously described^22^ and by NLuc bioluminescence assay of faecal output.

#### 2.5.1 Propagation of reporter parasite strains

Oocysts from the fourth passages of the FS3N strain in mice were purified and used to initiate passage in calves (**Supplemental Figure 2A**). For each passage (**Supplemental Table 2**), oocysts were prepared in PBS (Thermofisher). Cattle do not generally require oral gavage for this purpose so were directly given oocysts, delivered orally in 10 ml PBS.

For passage in calves, animals were euthanised once infection began to resolve; informed by NLuc results (**Supplemental Figure 2C**). Oocysts were purified from faecal material pooled from all infected animals from timepoints with the highest level of shedding.

#### 2.5.2 Calf Tissue Atlas

Study design was approved by the Moredun Research Institute Animal Welfare Ethical Review Board (AWERB), approving the use of the minimum number of animals per group (*n* = 4 per group).

Eight male neonatal calves (**Table 1**) were purchased at birth from two farms local to Moredun Research Institute. All animals passed a visual health inspection upon arrival at Moredun to satisfy enrolment criteria. The first four calves to arrive were assigned to Group 1 and were designated for experimental infection (“Infected” group). The second four calves were then assigned to Group 2 and were designated as uninfected controls (“Control” group). Group 1 were fitted with a harness and bag apparatus for faecal output collection throughout the study.

**Table 1.** Description of animals enrolled in study. Group assignment, breed, identification number, date of birth (corresponding to individuals reported in **Figure 1**).

| **Group** | **Calf Number** | **Breed** | **Date of Birth** |
| --- | --- | --- | --- |
| Infected | 1 | Jersey Cross | 17/04/24 |
|  | 2 | Holstein Friesian | 17/04/24 |
|  | 3 | British Blue Cross | 17/04/24 |
|  | 4 | British Blue Cross | 17/04/24 |
| Control | 1 | Holstein Friesian | 21/04/24 |
|  | 2 | Holstein Friesian | 21/04/24 |
|  | 3 | Holstein Friesian | 21/04/24 |
|  | 4 | Holstein Friesian | 21/04/24 |

As part of the enrolment process, faecal samples from all eight animals were tested for pathogens commonly implicated in cattle scours: Rotavirus, bovine Coronavirus*, E. coli* F5 (K99), *Cryptosporidium parvum*, *Clostridium perfringens,* using Rainbow™ Test Kits (Biox Diagnostics, BIO K 306). All eight animals tested positive for *C. perfringens* but negative for other pathogens tested.

##### i. Infection

All animals were housed in Moredun’s High Security Unit, in specific pathogen-free rooms. Group 1 animals were housed at a higher containment level than Group 2 due to infection with transgenic parasites. Group 1 animals were individually penned. On Day 0 (“Start”), Group 1 animals received 7.5×10^6^ FS3N oocysts each, administered in 10 ml PBS. These oocysts were purified from the second calf passage. Group 2 animals were not mock infected.

Infection levels were monitored until all Group 1 calves reached similarly high levels of infection, at which point (day 8 post infection) they were euthanised for tissue collection (**Supplemental Table 3**). After the Group 1 cohort was infected, calves for Group 2 were recruited. Staggering the two groups was critical to facilitate intensive post-mortem tissue collection. Group 2 animals were likewise euthanised eight days after their experimental start date to facilitate comparison to Group 1.

Animals’ feeding habits (**Supplemental Table 4**) and demeanour (**Supplemental Table 5**) were assessed twice daily as part of their welfare checks. Faecal output was monitored at least twice daily, and bags changed if containing faeces. Contents of bags were examined and allocated a score of 1-4 based on their consistency/diarrhoeal severity, where 1 = normal consistency and 4 = severe watery diarrhoea/scour. Scoring was completed by the same individual for all samples. From each animal’s faecal output, infection levels were individually quantified by NLuc assay.

##### ii. Tissue Collection

Ten locations along the GI tract were selected for sampling. The locations were chosen based on identifiable biological landmarks to ensure sampling areas were consistent between animals. Tissues were also collected in the same order for each animal (**Table 2**). Approximately 10 cm tissue sections were removed from each location. Tissue was cut longitudinally to open it and further divided into 1 cm x 1 cm pieces for various downstream analyses. Gridded DispoCut™ A4 dissection boards (CellPath) were used to collect each tissue sample individually to ensure samples of consistent size and eliminate contamination between sampling locations.

**Table 2.** Description of calf tissue sampling. Samples collected in this order. Corresponding regions collected from mouse tissue.

| Sample | Tissue | How tissue was identified |
| --- | --- | --- |
| 1 | Duodenum | ~10cm from abomasum |
| 2 | Jejunum B | Jejunum coils. Sample taken after 2 coils in from the proximal end |
| 3 | Jejunum E | Jejunum coils. Sample taken from 2 coils away from the distal end |
| 4 | Jejunum M | Approximately halfway along the jejunum |
| 5 | Ileum B | Entire ~30cm section removed and divided into three parts. Identified by proximity to the caecum |
| 6 | Ileum M |  |
| 7 | Ileum E |  |
| 8 | Colon | Ascending; closest to the caecum |
| 9 | Ileo-caecal Junction (ICJ) | Junction between caecum and ileum |
| 10 | Caecum | Tip of Caecum's blunt end/furthest point from ICJ |

Oesophageal tissue was similarly collected from individual “Control 1” from Group 2 to use in the generation of the standard curve for parasite quantitation from tissue by qPCR (as described in **2.8**).

### 2.6 Oocyst Purification

Oocysts were purified from mouse faecal samples via sucrose floatation and caesium chloride density gradient as previously described^19^. Oocysts were purified from calf faecal samples via acid flocculation as previously described^23^.

### 2.7 Parasite Quantitation by NanoLuciferase Assay

20 mg of faecal sample, or 20 mg of finely cut tissue were weighed into a 1.5 ml microcentrifuge tube with ∼10 glass bashing beads and homogenised in 1 ml of faecal lysis buffer^19^. Samples were agitated at room temperature at 2,000 rpm for 3 minutes (faeces) or 15 minutes (tissue) using an Eppendorf ThermoMixer C (#5382000031). 25 µl of a 1:50 ratio of Nano-Glo (Promega) assay buffer: substrate was combined with 100 µl homogenised sample in an opaque white plate without a lid (VWR, # PERK6005290). Samples were prepared in triplicate. Luminescence was measured using a Promega GloMax Navigator GM2000 (mouse samples, Dundee) or Promega GloMax Multi+ Detection System plate reader (calf faecal samples, Moredun Research Institute).

### 2.8 Parasite Quantitation from tissue by qPCR

Standard curve was prepared by extracting DNA from 25 mg of calf oesophageal tissue (collected from individual “Control 1” from Group 2) mixed with a specified number of wild type oocysts. 25mg of finely cut tissue was weighed into a 1.5 ml microcentrifuge tube prior to DNA extraction using DNeasy Blood & Tissue Kit (Qiagen). Samples were freeze-thawed five times in liquid nitrogen after the first step of the kit’s protocol. DNA was eluted in 25 µl of pre-warmed (50-60 °C) elution buffer and was eluted twice through the column. Extracted DNA samples were stored at −20 °C prior to qPCR. *C. parvum* was quantified as previously described^24^.(**Supplemental Table 1**). PCR was performed using Luna Universal Probe qPCR Master Mix (NEB) and a QuantStudio 3 qPCR System (Applied Biosystems) (**Supplemental Figure 3**).

### 2.9 *Cpgp*60 subtyping

Control animal tissue samples that were qPCR-positive for *C. parvum* were subtyped for strain identification. A nested PCR targeting the *C. parvum gp60* gene (**Supplemental Table 1**) followed by Sanger sequencing was used to subtype samples as described^25^. 5 µl of DNA that was extracted for qPCR (see **2.9**) was used as the target for the primary PCR (external PCR). A 20 µl reaction mix was prepared for each sample, using PrimeSTAR Max 2X Master mix (Takara Bio, #R045). The product of the primary PCR (external PCR) was diluted in 100 µl of dH_2_O, and 5 µl of this was used as the target for the secondary PCR (internal PCR). The resulting amplicon was purified via gel extraction (Monarch DNA Gel Extraction Kit, NEB) prior to sequencing via bi-directional Sanger sequencing. Analysis of sequences for subtyping was conducted manually as previously described^26^. Sanger Sequencing for *Cp*gp60 subtyping was conducted by the University of Dundee MRC PPU DNA Sequencing and Services.

### 2.10 Microbiome Analysis

On day 0 (study start, D0) and day 8 (study end, D8), faecal material from each calf was sampled for microbiome analysis. Approximately 150 mg of faeces were collected into barcoded tubes stored at room temperature. Samples were processed and analysed by Transnetyx (Cordova, Tennessee, USA). Sample preparation was conducted by Transnetyx. In brief, DNA was extracted using DNeasy 96 PowerSoil Pro QIAcube HT extraction kit (Qiagen). Library preparation was performed using the KAPA HyperPlus library preparation protocol (Roche) optimised for minimal bias. After QC, libraries were sequenced (Illumina NextSeq 2000) via shotgun sequencing at a depth of 2 million 2x 150 bp read pairs, giving species and strain level taxonomic resolution. Unique dual indexed (UDI) adapters were used to ensure that reads and/or organisms were not mis-assigned. Data was analysed by Transnetyx and provided via the One Codex database^27^ and interface.

### 2.11 Confocal Microscopy (live oocysts)

2 million oocysts were incubated with 2% bleach on ice for 5 minutes and washed 3x with PBS as previously described^19^. Oocysts were centrifuged at 16,000 x *g* at 4 °C for 3 minutes and resuspended in 20 µl Matrigel (Corning) before loading onto a µ-Slide Angiogenesis microscope slide (Ibidi, #81501). Oocysts were allowed to settle overnight at 4 °C. Following this overnight incubation, the slide was incubated at 37°C for 15 minutes to solidify the Matrigel before imaging. Imaging was performed at Dundee Imaging Facility using a Zeiss LSM880 Airyscan microscope (confocal mode).

### 2.12 Confocal Microscopy (Tissue)

#### i. Sample Preparation

Tissue sections (1cm x 1cm) were fixed overnight in 4% paraformaldehyde in PBS (Sigma catalogue #441244). Samples were washed 3x with PBS and then incubated overnight in 30 ml 30% sucrose at 4 °C. Samples were then briefly washed again with PBS and embedded in optimal cutting temperature (OCT) compound (Scigen, #4568). This was performed over a dry ice + 100% ethanol bath to freeze samples and solidify OCT whilst embedding. Blocks were stored at −20 °C prior to cryo-sectioning (Leica CM1850 cryostat). 12 μm sections were cut at - 24 °C and were dried onto slides overnight prior to staining.

To stain, slides were permeabilised with 1% Triton-X-100 at room temperature for 10 minutes and then washed 3x with PBS. Slides were stained for 30 mins with Hoechst 33342 (Thermofisher; 1:2000 in PBS) and Phalloidin-647(Abcam, #ab176759; 1:1000 in PBS), then washed 3x with PBS and mounted using ProLong™ Glass Antifade Mountant (ThermoFisher). Slides were cured at room temperature overnight prior to imaging.

#### ii. Imaging

Imaging was performed at Dundee Imaging Facility using a Leica Stellaris 8 confocal microscope. All images were captured using the 40x oil immersion objective (HC PL APO CS2 40x/1.30 OIL, Leica). The tuneable laser of the microscope was adjusted to the peak excitation wavelength for each fluorophore. All images were collected using the same laser settings, at pinhole 1. Laser settings were consistent between images (Hoechst: 31.7% gain, 2.00% intensity; mNeon: 112.4 gain, 4.35% intensity; Phalloidin: 10.0% gain, 1.08% intensity).

All z-stacks were collected at a step distance of 1 μm, encompassing the thickness of the slice. From z-stacks of Ileum and Ileo-caecal junction, z-projections were produced using Fiji ver. 1.53q (ImageJ). Channels were merged, and z-projections created using the max intensity z-project function. To reduce background, the colour balance of the Ileum z-projection was adjusted to reduce levels of white and green background, and the colour balance of the ICJ z-projection was adjusted to reduce levels of blue and green channel (Hoechst and mNeon) background. No other adjustments were made.

### 2.13 Scanning Electron Microscopy

Samples were prepared and images were collected at the Cellular Analysis Facility, College of Medical, Veterinary, and Life Sciences Shared Facilities Resources, University of Glasgow. Samples were prepared and imaged blind to identity of sample (infected or control). Upon collection, tissue sections (1cm x 1cm) were fixed in 10 ml total 4% paraformaldehyde (Sigma, #441244) with 2.5% glutaraldehyde (Merck, #G6257) and stored at 4 °C. To prepare for SEM, the tissue sections were washed in cacodylate 0.1M buffer, followed by dehydration in ascending series of ethanol (from 30% to 100%), and critical point dried using the Autosamdri815 system (Tousimis, USA). Dried samples were mounted on aluminium stubs covered with carbon tape and fixed in place with conductive silver paint (Agar Scientific). Afterwards, they were coated with gold/palladium (5 nm thick layer) and imaged using the Jeol IT-100 scanning electron microscope.

### 2.14 Statistical Analysis and Software

Graphs were produced using GraphPad Prism version 10.2.2. qPCR results were visualised using Design & Analysis (DA1) software (Fisher Scientific) and parasites/g tissue was calculated using Microsoft Excel and GraphPad Prism version 10.2.2. Microbiome data was visualised using TransnetYX/OneCodex Quick Compare tool. Confocal microscopy images were collected at the Dundee Imaging Facility; The Open Microscope Environment (OMERO)^28^ was utilized for image management (https://www.openmicroscopy.org/omero/). Confocal microscopy z-projections were generated using Fiji ver. 1.53q (ImageJ).

### 2.15 Power analysis

Power analysis was performed using data generated from the calf infection trial in this study, in order to be useful for determining group sizes in future animal trials. Anticipated means and standard error were based on the mean NLuc activity from tissue and faecal samples at peak shedding in *C. parvum*-infected calves (**Supplemental Table 6**). The anticipated mean for a treatment group (in a two independent study groups model) was based on either a 90% or 99% reduction in NLuc activity. Power was calculated at 80% (β = 0.2) and 90% (β = 0.1), respectively. Alpha was set to 0.05 and an enrolment ratio of 1 was used.

## 3. Results

### 3.1 An infection model for moderate cryptosporidiosis in neonatal calves using transgenic reporter parasites

To test the hypothesis that infection of the calf gut with *C. parvum* is restricted to the ileo-caecal junction (ICJ), we performed a first-of-its-kind infection study to map and quantify *C. parvum* infection levels throughout the neonatal calf gastrointestinal (GI) tract. We aimed to analyse this in the context of a moderate cryptosporidiosis infection. Moderate infection is most often observed in agricultural settings, and although not fatal it still causes animal morbidity and has an economic impact^4^.

To improve ease of detection and quantitation of *Cryptosporidium* infection, we utilised a transgenic *C. parvum* reporter strain. Expression of NanoLuciferase (NLuc) allows very sensitive quantitation of parasites in tissue and faecal samples, expression of fluorescent proteins enables high-resolution microscopy of infected tissue, and paromomycin resistance allows for selection (**Supplemental Figure 1**). The reporter strain was generated and propagated in IFNγKO mice and then passaged twice in neonatal calves prior to this experimental study (**Supplemental Figure 2** and **Supplemental Table 2**).

Eight calves were enrolled at birth; the first four were allocated to the “Infected” group and the next four to the “Control” group (**Table 1** and **Figure 1**). Upon enrolment in the study, all calves were treated with paromomycin. In addition to selecting for growth of the reporter strain, treatment with paromomycin reduces the possibility of infection with wild type parasites that calves may be exposed to at birth, especially important for the “Control” group. Calves in the “Infected” group were orally infected with reporter parasites on DPI 0 (day post infection; **Figure 1A**).

**Figure 1.**
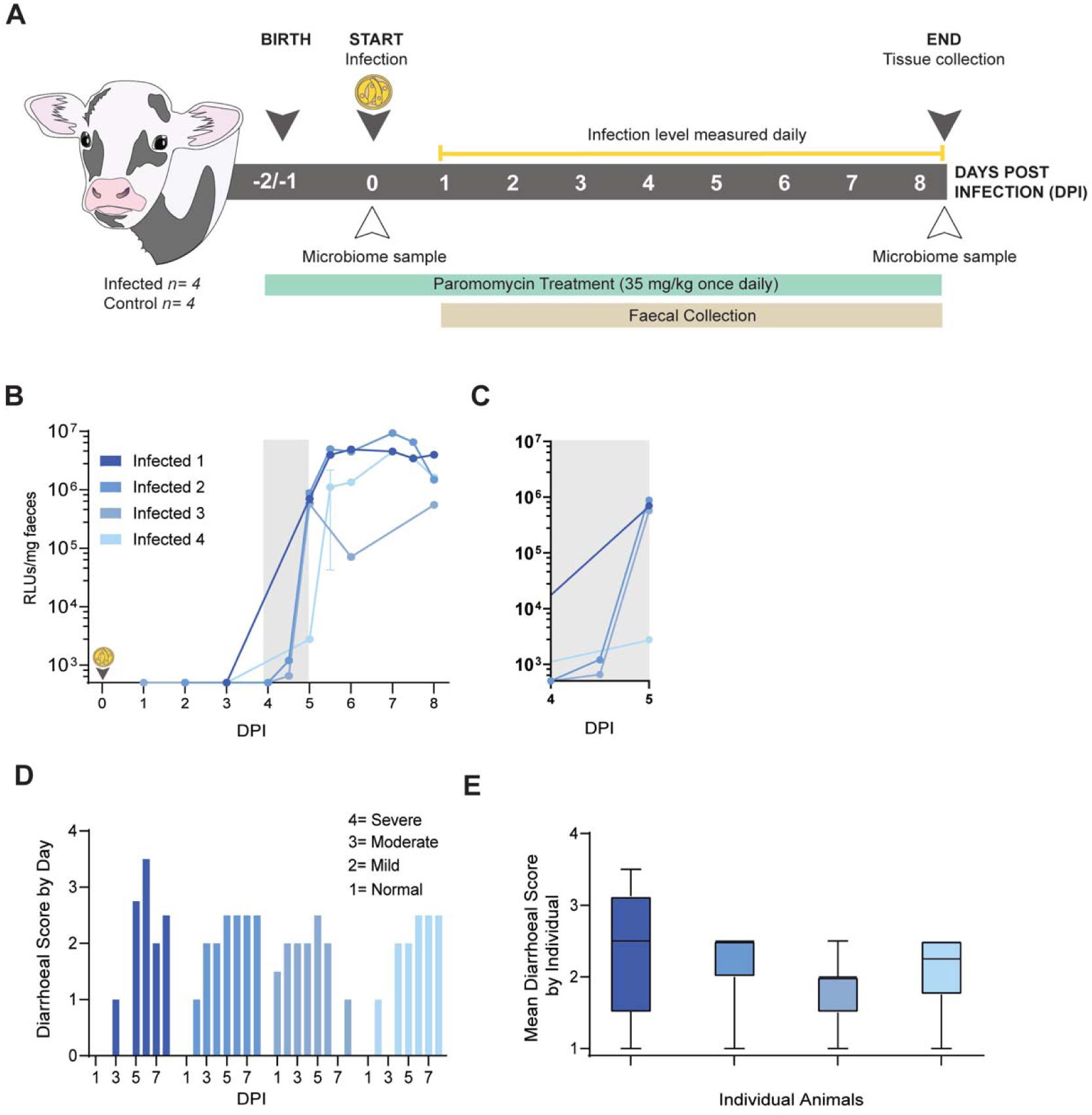
Acute experimental cryptosporidiosis of neonatal calves produces a moderate infection. **A**) Schematic of neonatal calf infection study (grey box). Eight male calves were recruited at birth (grey arrow), with four calves assigned to the experimental “Infected” group and four calves to the uninfected “Control” group. All calves received paromomycin throughout (green box). The study began on day post infection (DPI) 0 with oral infection of calves with transgenic *Cryptosporidium parvum* (indicated by yellow oocyst). Faecal samples were collected (brown box) throughout the study and analysed for *Cryptosporidium* shedding (yellow line). The study ended on DPI 8, when animals were euthanised and GI tract tissue collected. Microbiome of faecal samples was analysed on DPI 0 and 8 (white arrows). All calves received paromomycin throughout (green box). The transgenic strain of *C. parvum* used in this study is resistant to treatment with paromomycin sulphate. **B**) Parasite shedding of individual animals (blue) from the infected group was quantified via faecal NanoLuciferase assay (RLUs/mg faeces). Where calves produced 2 samples a day, both are plotted; where 3 samples were produced the second and third were averaged for the second data point (DPI 5, Infected 4). Grey box indicates time of observed exponential increase in shedding. Average ± SD plotted; 3 or 6 technical replicates plotted for each sample. Y-axis set at limit of detection (500 RLUs/mg). **C**) Inset of (**B**). **D**) Mean daily diarrhoeal score (mean of 1-3 scores per day, per individual). Where no bar is present, no sample was produced to be scored. **E**) Diarrhoeal score was not significantly different between individual infected animals (unpaired, two-tailed *t*-test, *p* > 0.05 for all). Diarrhoeal score for individuals over the course of the study (centre is median of all scores from DPI 0 to 8; whiskers denote minimum and maximum scores).

Parasite shedding in faecal samples was monitored for “Infected” individuals via faecal NLuc assay (**Figure 1B**). Neonatal calf cryptosporidiosis is acute, with peak shedding typically occurring approximately one-week post-infection and resolving two weeks post-infection. All “Infected” individuals reached high levels of parasite shedding, with shedding for three of four individuals increasing exponentially DPI 4-5 to >5.5 x 10^5^ RLUs/mg (**Figure 1C**). Peak NLuc activity in faecal samples peaked between DPI 5-8, with a slight reduction by DPI 8 in two of the infected calves, suggesting oocyst shedding had peaked. Therefore, animals were culled at DPI 8 (1.91×10^6^ ± 1.45×10^6^ RLUs/mg mean and SD of individuals at DPI 8; individual faecal NLucs reported in **Supplemental Table 3**).

Total faecal output was collected daily from each “Infected” individual, and diarrhoea was scored (**Figure 1D**). While calves became highly infected and shed large quantities of *C. parvum*, each individual experienced mild-to-moderate diarrhoeal scores throughout, and no individual produced severely diarrhoeic samples. Diarrhoeal scores were consistent between individuals (**Figure 1E**), with no significant difference between each individual’s mean diarrhoeal scores. Animals’ welfare and behaviour was assessed with twice daily scoring for feeding and demeanour (**Supplemental Tables 4 and 5**). There was no difference observed between “Infected” and “Control” groups. Other than diarrhoeal symptoms, “Infected” animals appeared otherwise healthy. This confirms that infected animals experienced moderate cryptosporidiosis, where large numbers of parasites were shed and diarrhoea was observed, but without acute impact on welfare.

### 3.2 Tissue atlas of *Cryptosporidium* infection in neonatal calves reveals infection throughout the gastrointestinal tract

Once “Infected” individual calves achieved peak shedding, they were euthanised for tissue collection on DPI 8. Samples were collected from each calf for both “Infected” and “Control” groups. Tissues were collected from regions that could be reproducibly sampled from individual to individual and located throughout the GI tract (Figure **2A** and **C** and **Table 2**). Infection was quantified by NLuc assay (**Figure 2D**) and by qPCR (**Supplemental Figure 3**). All calves became well infected, and we observed similar levels of infection of the tissues from individual-to-individual.

**Figure 2.**
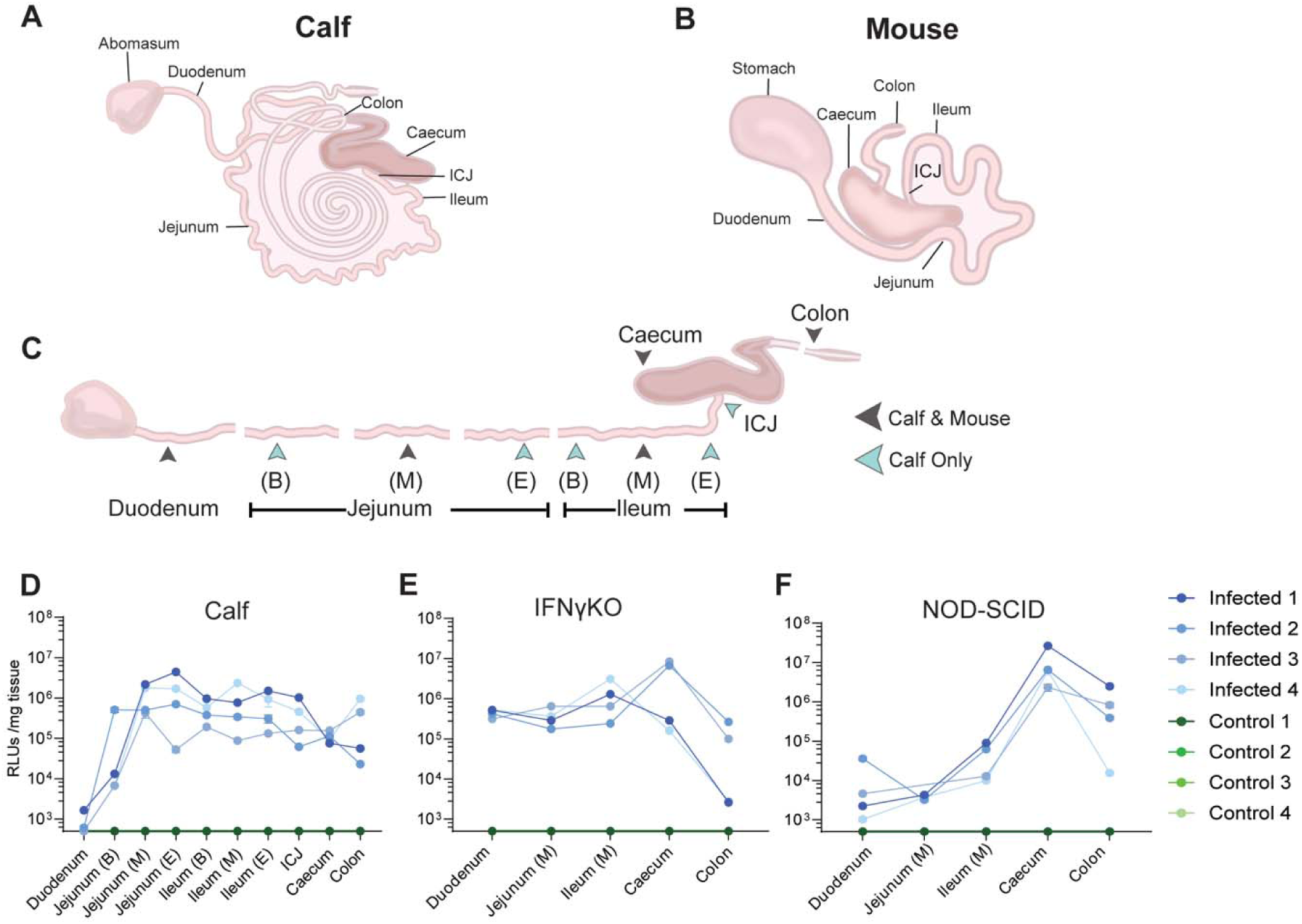
Tissue atlas of *Cryptosporidium parvum* infection models. GI tract of experimentally infected animal models, neonatal calf and immunocompromised mouse, were sampled to quantify parasite burden. **A**) Anatomy of neonatal calf GI or (**B**) mouse with **C**) sampling locations indicated. Diagrams for illustrative purposes and not to scale. ICJ, ileo-caecal junction. Ten locations were sampled for calves (all; **Table 2**). Due to the length of the jejunum and ileum, these regions were sampled at the beginning (‘B’), middle (‘M’), and end (‘E’). Duodenum was sampled close to the abomasum and colon sampled near the caecum. and five sample locations for mice (green arrow). For mice, five corresponding sampling locations (grey arrows) were collected to enable direct comparison. Fewer samples were collected overall, with jejunum and ileum sampled only in the middle (‘M’). **D-F**) Parasite burden in GI tract tissue of infected individuals (blue) and control animals (green) was quantified via NLuc assay (RLUs/mg tissue). Mean ± SD; mean of 3 technical replicates plotted. Y-axis set at limit of detection (500 RLUs/mg). Data points less than or equal to the limit of detection were plotted as such.

Counter to previous reports, we observed that infection with *C. parvum* is not limited to the ileo-caecal junction but is present at high levels throughout the GI tract (**Figure 2D**). Within an individual, infection level was relatively consistent throughout the small intestine (in the range of 10^5^ to 10^7^ parasites/gram tissue; **Supplemental Figure 3**), except for lower levels at the duodenum and beginning of the jejunum (<10^5^ parasites/gram tissue). For each individual, the caecum was consistently infected at lower levels than infection in the small intestine. Infection levels were especially low in the duodenum, with levels at or only marginally above the level of detection by NLuc and below the limit of quantitation by qPCR.

As expected, tissue samples from the “Control” group did not have detectable levels of *C. parvum*, as determined by NLuc assay. By qPCR, “Control” individuals were negative except for two samples (the beginning of the ileum in Individual 1 and beginning of the jejunum in Individual 4; **Supplemental Figure 3**). These samples were above the limit of quantitation, but very lowly infected (2.52 x 10^4^ parasites/g tissue and 5.48 x 10^3^ parasites/g tissue respectively), relative to the infection level recorded in the “Infected” group at these tissue sampling sites. We determined these samples to be *Cpgp60* IIaA17G2R1 subtype. This is the same *gp60* subtype as that of the reporter strain used and is prevalent in the geographic region where the calves were sourced. Therefore, we attribute these two lowly positive samples to exposure of the “Control” calves to *C. parvum* in the farm environment before enrolment in the study. Treatment with paromomycin was successful in reducing this environmental exposure as *Cryptosporidium* was “undetectable” in all other “Control” samples.

### 3.3 Tissue Atlas of cryptosporidiosis immunocompromised mouse models reveals important differences in infection patterns

Wild type mice are not well infected with *C. parvum*^24^. Therefore, immunocompromised mouse models are commonly employed in laboratory settings to model *C. parvum* infections of the natural host, in this instance neonatal calves. To understand how well these immunocompromised mouse models recapitulate the infection pattern observed in the neonatal calf GI tract, we generated a tissue atlas for both the acute (IFNγKO mice) and chronic (NOD-SCID gamma, NSG mice) cryptosporidiosis mouse models to compare them to the calf tissue atlas.

Both mouse models were infected with the reporter strain used in the calf tissue atlas study for this purpose. Mice were culled at peak infection (**Supplemental Table 3**) and tissue samples were collected (**Figure 2B and C**) in regions corresponding to those sampled for the calf tissue atlas (**Table 2**). Infection levels in the tissue were quantified via NLuc assay.

In IFNγKO mice, infection is widespread throughout the small intestine (**Figure 2E**). Infection levels are similar at various sites sampled in the small intestine. This includes high levels of infection in the duodenum (4.46×10^5^ ± 8.67×10^4^ RLUs/mg, mean and SD of four mice). This is markedly different than observations from the calf tissue atlas (8.18×10^2^ ± 5.19×10^2^ RLUs/mg, mean and SD of four calves). Although the colon of IFNγKO mice was positive for *C. parvum*, infection was observed to be the lowest here (9.26×10^4^ ± 1.12×10^5^ RLUs/mg, mean and SD of four mice; 3.68×10^5^ ± 3.91×10^5^ RLUs/mg, mean and SD of four calves). Other than the infection of the duodenum, the tissue infection pattern of neonatal calves and IFNγKO mice was observed to be very similar.

In contrast to the infection profile across GI tract tissue sites in the bovine calf and IFNyKO mice, in the NSG mouse *C. parvum* is present at low levels in the small intestine (4.37x 10^4^ ± 3.54×10^4^ RLUs/mg, mean and SD of four mice ileum sample; **Figure 2F**). Infection levels increase further along the length of the GI tract, with the highest level of infection in the caecum (NSG mice 1.04×10^7^ ± 9.94×10^6^ RLUs/mg, mean and SD of four mice; IFNγKO mice 3.9×10^6^ ± 3.89×10^6^ RLUs/mg, mean and SD of 4 mice; calves 1.09×10^5^ ± 3.10×10^4^ RLUs/mg, mean of four calves) and the second highest level of infection observed in the colon.

Additionally, in separate experiments, NSG mice were infected with a reporter strain^20^ of *C. parvum* that expresses NLuc and a red-shifted firefly luciferase (FLuc) to enable both *in vivo* and *ex vivo* imaging. A time course of *in vivo* imaging of infected NSG mice confirms chronic infection (**Figure 3 A-C**). *Ex vivo* imaging of GI tract tissue of chronically infected NSG mice confirms the highest infection levels occur in the caecum and colon irrespective of animal supplier (**Figure 3D**). This infection is significantly different from the tissue distribution observed for neonatal calves.

**Figure 3.**
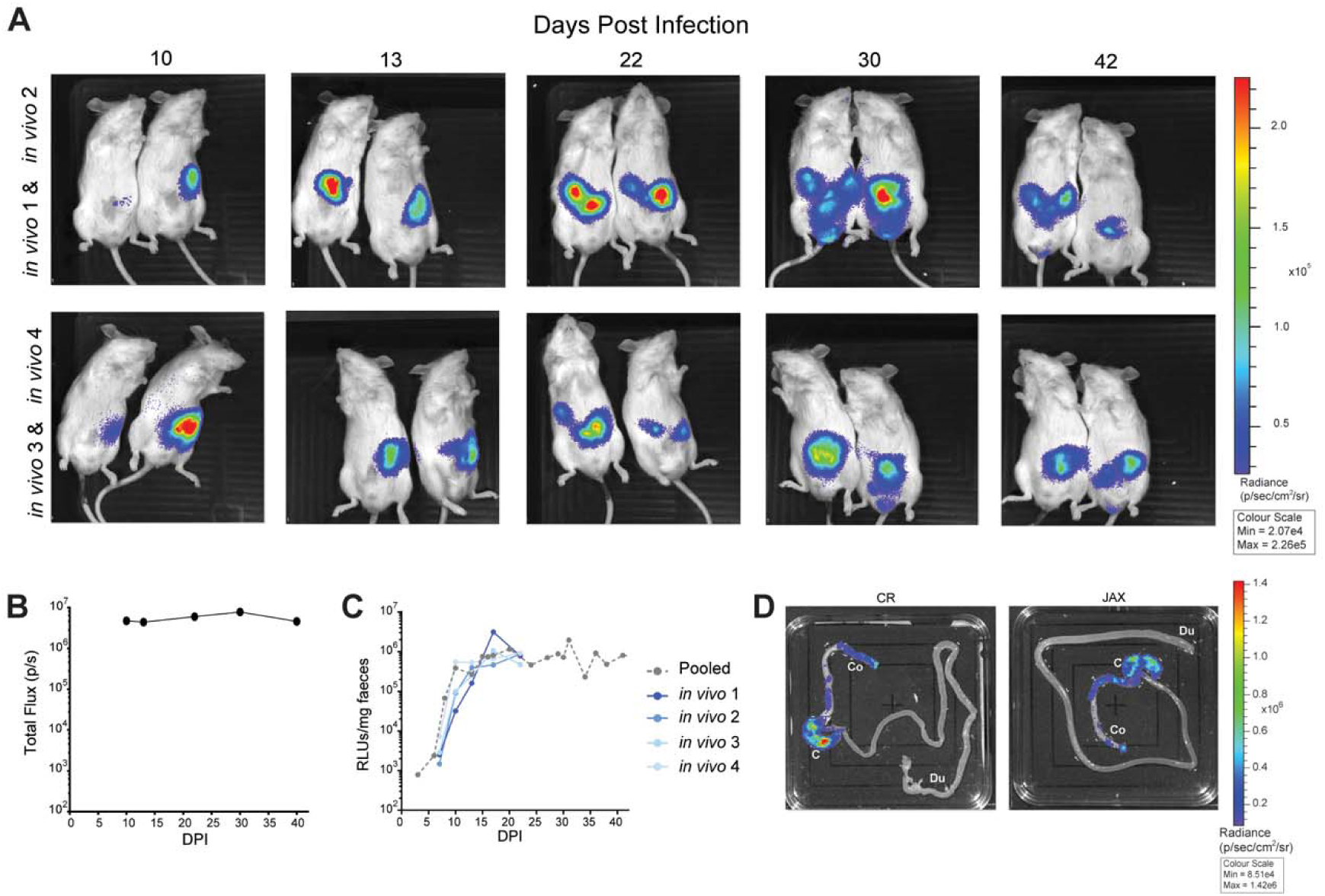
*In vivo* and *ex vivo* imaging of chronic cryptosporidiosis mouse model, NSG mice. **A**) *In vivo* imaging of chronic cryptosporidiosis post infection **B**) Infection quantified as total flux (photons/sec; mean ± SD of *n*=4 mice plotted). *In vivo* infection levels correspond to **C**) shedding of parasites in faecal samples (RLUs/mg faeces; mean ± SD; mean of 3 technical replicates plotted. Y-axis set at limit of detection of 500 RLUs/mg). **D**) *Ex vivo* imaging of GI tract from mice culled at peak infection (animals purchased from different suppliers: “CR” Charles River, UK and “JAX” Jackson Labs, USA). Tissue locations of interest labelled: duodenum, Du; caecum, C; colon, Co. Representative image shown; *ex vivo* tissue samples from additional individuals in **Supplemental Figure 4**.

### 3.4 Visualisation of *Cryptosporidium* parasites in the neonatal calf GI

*Cryptosporidium parvum* infection from throughout the GI tractof “Infected” neonatal calves was visualised using confocal microscopy (**Figure 4**). *C. parvum* (expressing cytoplasmic mNeon) were observed in all GI tract samples, with observably lower levels in duodenal and caecal tissue compared to others (**Figure 4A**). Parasites heavily infect the brush border of gut epithelial cells. In particular, the ileum (**Figure 4B**) and ICJ (**Figure 4C**) tissue showcase noticeably higher levels of infection, with parasites populating a large portion of the villus surface area.

**Figure 4.**
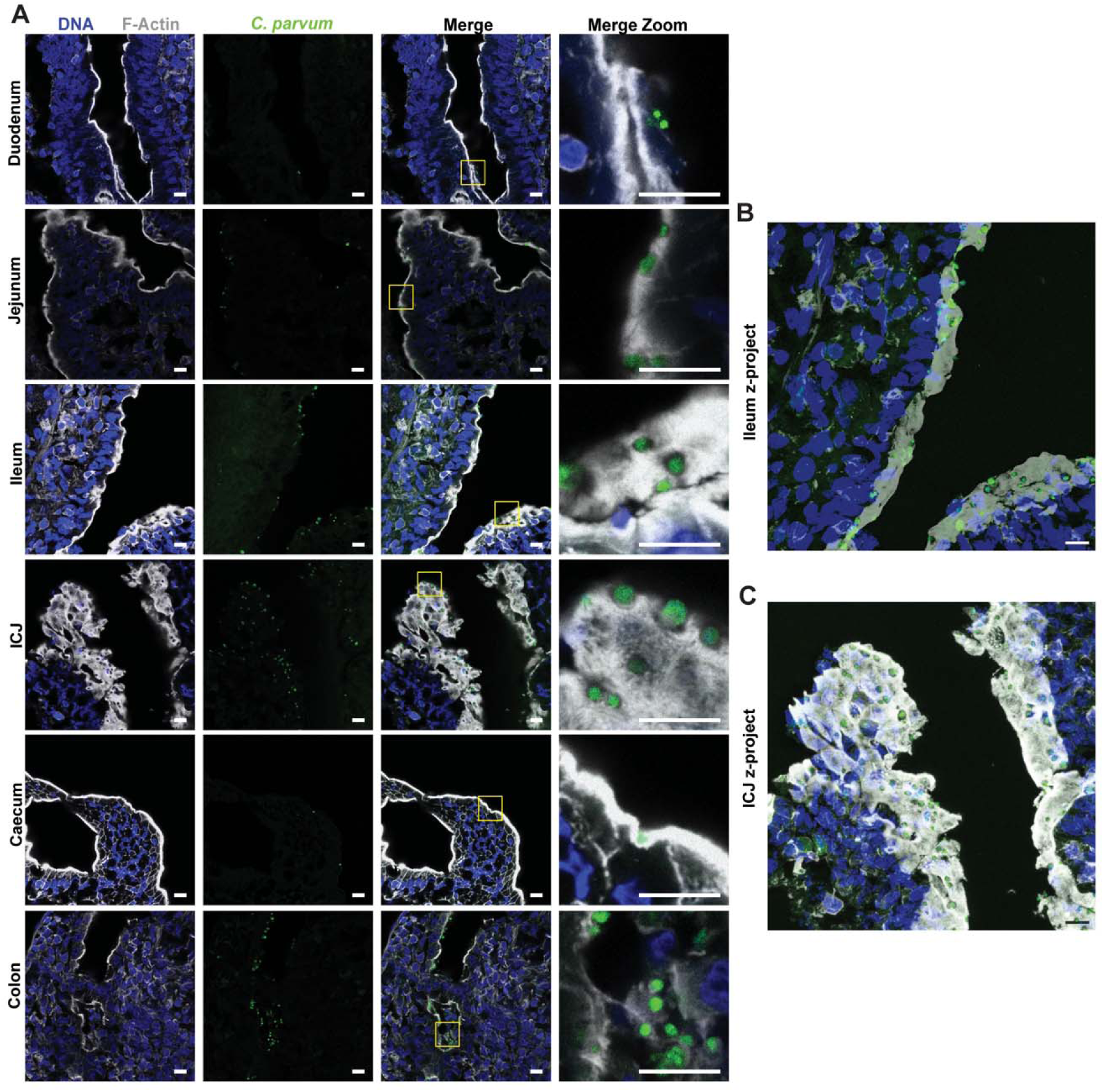
*C. parvum* infection visualised throughout the neonatal calf GI. Tissue samples collected throughout the GI tract (duodenum, middle of jejunum, middle of ileum, ileo-caecal junction (ICJ), caecum, colon) of an individual infected calf (“Infected 1”, see Figure 2D) were imaged using confocal microscopy (**A**, single z-plane) and prepared as maximum intensity projections of the ileum and ICJ (**B** and **C**, several z-stacks). Gut tissue morphology (brush bordre) visualised using a stain for F-actin (white, Phalloidin-647) and DNA (blue, Hoechst). Reporter transgenic *C. parvum* strain visualised via cytoplasmic mNeon (green). “Merge zoom” column corresponds to inset of “Merge” column (yellow box). Scale bar 10 µm for all images. Images were collected using the 40x oil immersion objective (HC PL APO CS2 40x/1.30 OIL, Leica), with the tuneable laser adjusted to the peak excitation wavelength for each fluorophore. All images were collected at pinhole 1 AU. Laser settings were consistent between images (For Hoechst: 31.7% gain, 2.00% intensity; mNeon: 112.4% gain, 4.35% intensity; Phalloidin: 10.0% gain, 1.08% intensity). Z-projections were generated as a projection of 38 and 55 z-sections (**B** and **C**, respectively).

High levels of infection in the ileum were further observed using scanning electron microscopy (SEM; **Figure 5A and B**). Ileal tissue from each calf was visibly studded with large numbers of parasites. Here, areas of infection are clearly visible, and in all infected animals, parasites are visibly surrounded by elongated microvilli. SEM of control animal ileal tissue (**Supplemental Figure 5A**) shows normal ileum morphology. We identified a single, rare infection event (**Supplemental Figure 5B**) in the ileal tissue of a control calf, corresponding to a sample that was reported by qPCR (**Supplemental Figure 3)**.

**Figure 5.**
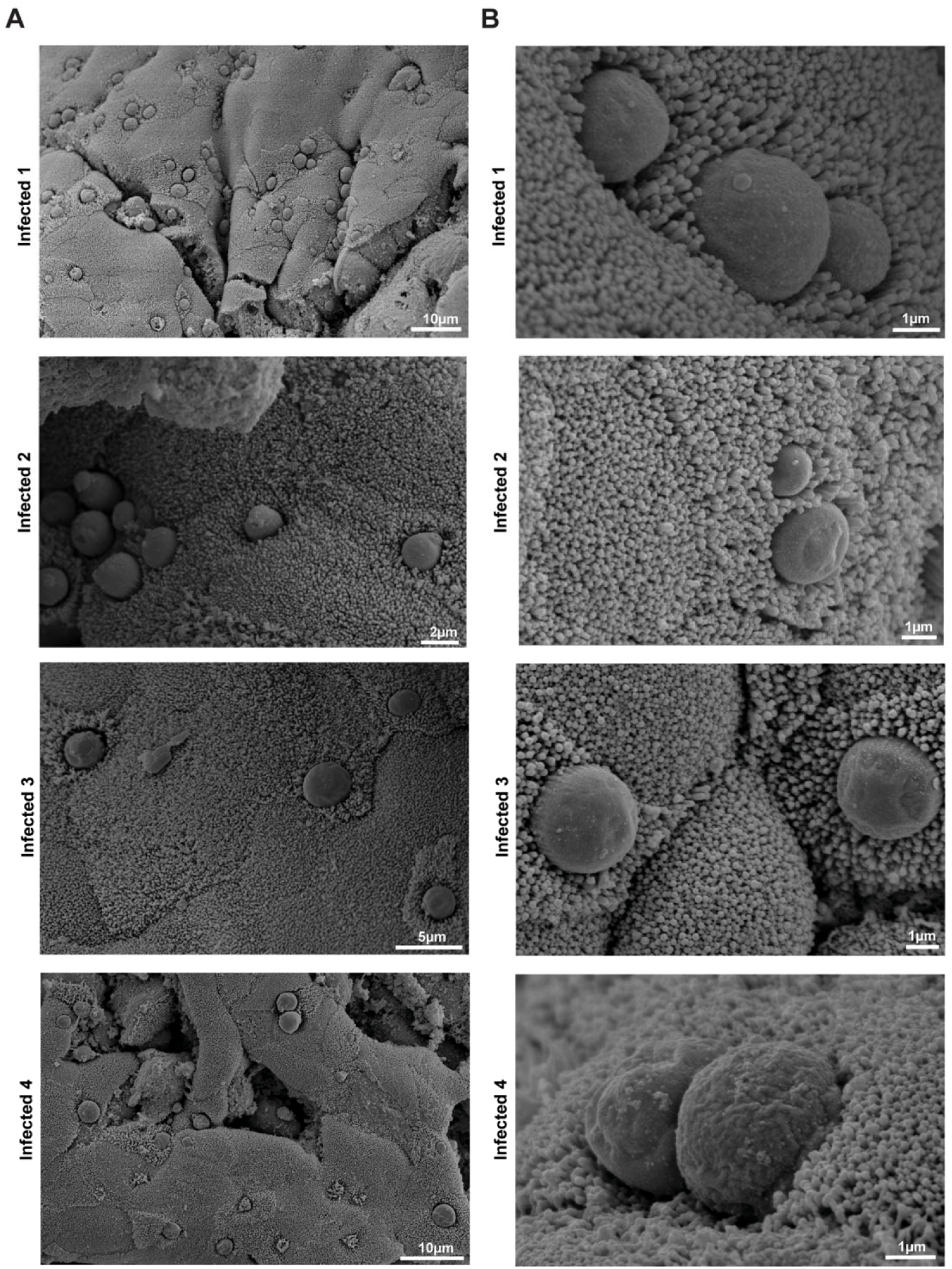
Scanning Electron Microscopy of infected calf ileum. Tissue samples collected from the middle of the ileum of all infected calves (Infected 1 - 4) were imaged using scanning electron microscopy (**A** lower magnification, **B** higher magnification). Corresponding tissue samples from control animals reported in **Supplemental Figure 5**.

### 3.5 Power analysis to inform future studies of moderate cryptosporidiosis in the neonatal calf model

Data from this first-of-its-kind study can be used as a guide for determining group sizes for future investigations that require neonatal calf infection models. This will be particularly useful for studies aiming to quantify reductions in *C. parvum* infection, namely drug efficacy studies.

Mean NLuc activity in *C. parvum*-infected tissue and faecal samples (± standard error) was used to calculate necessary animal group sizes, based on a reduction of 90% or 99% in infection. From this, estimated group sizes ranged from 1 calf per group to 23 calves per group, depending on the sample type (**Table 3**). Overall, duodenal and proximal jejunal tissue sites were less infected and thus less reliable for detecting a drop in *C. parvum* infection, due to the relatively lower infection burden at these sites in *C. parvum*-infected calves. Sampling sites at, or adjacent to, the ileo-caecal junction, and Day 6 and Day 8 faecal samples, were found to be more robust sampling sites, necessitating lower animal group sizes (**Table 3**).

**Table 3.** Power analysis calculation. Estimated group sizes necessary to detect a 90% or 99% reduction in *C. parvum* infection, based on power analysis using mean NLuc data measured from tissue or faecal samples as indicated (**Supplemental Table 6**).

| Tissue/Sample | Power 0.8 ($\beta = 0.2$) | | Power 0.9 ($\beta = 0.1$) | |
| --- | --- | --- | --- | --- |
|  | 90% reduction<br>(calves per group) | 99% reduction<br>(calves per group) | 90% reduction<br>(calves per group) | 99% reduction<br>(calves per group) |
| Duodenum | 9 | 8 | 12 | 10 |
| Jejunum (B) | 16 | 14 | 23 | 19 |
| Jejunum (M) | 3 | 2 | 3 | 3 |
| Jejunum (E) | 6 | 5 | 8 | 7 |
| Ileum (B) | 2 | 2 | 3 | 2 |
| Ileum (M) | 6 | 5 | 9 | 7 |
| Ileum (E) | 4 | 3 | 5 | 4 |
| ICJ | 5 | 4 | 7 | 6 |
| Caecum | 1 | 1 | 1 | 1 |
| Colon | 7 | 5 | 9 | 7 |
| Faeces (D6) | 4 | 3 | 5 | 4 |
| Faeces (D8) | 3 | 2 | 4 | 3 |

## 4. Discussion

To-date, there exist key gaps in our knowledge of *C. parvum* infection of natural hosts. Despite its use as research tool, the infection biology of cryptosporidiosis of neonatal calves has scarcely been investigated. The most foundational knowledge, namely the location and breadth of infection in the GI tract of this model, has yet to be robustly investigated. Currently our understanding of host-parasite biology is largely amassed from studies using immunocompromised mice. However, it has been unknown how well these laboratory immunocompromised mouse models (IFNγKO and NSG mice) recapitulate infection in the calf. This incomplete perspective hinders *in vivo* studies and drug discovery efforts. In this first-of-its-kind study, we created a tissue atlas of *C. parvum* infection of three important *in vivo* models, allowing new, robust comparisons to be drawn. This foundational knowledge provides new perspectives that will influence the way we study host-pathogen interaction.

We utilised and exploited transgenic parasites to facilitate our study. Since its introduction, CRISPR editing of *C. parvum* has revolutionised our ability to investigate the biology of this important parasite^17^. Our NLuc-expressing strain enables a simple, rapid, yet high-sensitivity readout of infection, both in faecal and tissue samples. Additionally, our strain is resistant to paromomycin, an approved anti-cryptosporidial for treatment in cattle. This was critical in reducing infection levels in “control” animals that may have been exposed to parasites from the farm environment. Our observation of low levels of *C. parvum* in two samples from two control calves highlights the importance of selection with paromomycin throughout the trial to prevent confounding observations between our experimental infection and wild type infection. We generated and passaged our reporter strain in IFNγKO mice and then passaged twice in calf models before conducting the tissue atlas study. To our knowledge, this provides the first reported serial passage of transgenic *C. parvum* in the neonatal calf model. The use of transgenic parasites was critical to the study’s success and development of the tissue atlas.

It is now clear that the true breadth of *C. parvum* infection in the neonatal calf GI tract was grossly underappreciated. These results challenge the long-standing hypothesis that infection is specific to one location (the ileo-caecal junction) and instead showcase the true widespread nature of parasite colonisation throughout the gut, with high levels of indiscriminate infection from the jejunum through to the colon. This provides reasoning as to why infected calves shed such high quantity of parasites. We also observe elongation of microvilli of epithelial cells infected with *Cryptosporidium*. This phenomenon was visualised previously in a bovine *Cryptosporidium sp.* infection^29^ and is recapitulated in severe combined immunodeficiency disease (SCID) mice infected with *C. parvum*^30^. Recent work in human intestinal epithelial cells and IFNγKO mice^31^ indicates the involvement of *Cryptosporidium parvum* Microvilli Protein 1 (*Cp*MVP1) and a family of related proteins that localise to the microvilli of the host cell and promote their elongation through interaction with a host scaffold protein. With our SEM images, we clearly visualise that this phenotype occurs during bovine infection as well.

These findings also present new considerations when designing and interpreting drug efficacy studies. Firstly, the distribution of *C. parvum* throughout the GI tract and the time course of infection observed in IFNγKO mice is very similar to neonatal calves. Therefore, this model may be more appropriate for shortlisting candidates for testing in calf models. In contrast, the distribution of parasites and the time course of infection is markedly different between neonatal calves and NSG mice. The NSG mouse may therefore be more appropriate when modelling chronic infection time courses and less effective for measuring drug efficacy. This correlation between efficacy testing in different host models should become apparent as candidate compounds progress to clinical development.

For animal health, it will be essential that anti-cryptosporidial drugs are present along the entire length of the GI tract to ensure all sites of infection are targeted. It is necessary to consider the pharmacokinetic-pharmacodynamic properties and dosage that will be sufficient to reduce parasite shedding in this model^32,33^. The results also provide a basis for power analysis to determine the experimental group size for future studies. From an ethical and logistical perspective, this data will be valuable for limiting the number of animals required for future experimental infection trials. Complete parasite killing is essential as recrudescence of disease can occur quickly and from a very low level of infection (below the limit of detection of oocysts in faecal samples). In addition, neonatal calves carry a higher burden of infection than humans, and emergence of parasite resistance has been observed^12^. As we now appreciate the large number of parasites that would be subjected to drug treatment, it will be important to consider the threat of drug resistance and the implications on transmission of drug-resistant parasites between animals and people.

We observe important differences between location of infection in the immunocompromised mouse models. In the NSG mouse, the burden of infection is several orders of magnitude higher in the caecum than elsewhere in the gut, showing a strikingly different infection pattern compared to both acute models. The underlying basis for the differences between each model remains unclear, however, it is possible that host-specific factors that vary between these hosts, such as immunological responses, or variations in physiological conditions (e.g. pH) might play a role. Further exploration of the underlying biology driving these changes will provide insight useful for better understanding of host-pathogen relationship.

As they are not the natural host and do not experience symptomatic diarrhoeal disease, immunocompromised mouse models are not the most relevant tool for understanding host-pathogen relationship. Our expanded understanding of the natural host will be useful in better modelling this relationship in the laboratory. New tools like calf-derived intestinal organoids for *in vitro* co-culture with *Cryptosporidium* could provide access to more physiological tools to study host-parasite interaction. There remains much to investigate regarding bovine cryptosporidiosis. The knowledge gained from this study as well as emerging tools for interrogating host-parasite interaction provide the foundation for important follow-up work. Integrating knowledge across human and animal health with a One Health approach is key to tackling this important parasitic disease.

## Supporting information

Goddard et al Supplemental

## Author Contributions

Peyton Goddard: Conceptualization, Methodology, Formal Analysis, Investigation, Writing-Original Draft, Visualization, Project Administration

Thomas Tzelos: Methodology, Investigation, Supervision

Beatrice L Colon: Conceptualization, Methodology, Formal Analysis, Investigation, Visualization, Project Administration

Paul M Bartley: Validation, Investigation

Lee Robinson: Conceptualization, Methodology, Formal Analysis, Investigation, Visualization Leandro Lemgruber: Investigation, Visualization

Michele Tinti: Methodology, Software, Formal Analysis, Visualization

Grant MJ Hall: Methodology, Validation

Sarah Stevens: Validation

Louise Gibbard: Investigation

Robert Bernard: Investigation

George Tytler: Investigation

David Smith: Conceptualization, Methodology, Formal Analysis, Resources, Writing-Review and Editing, Supervision, Project Administration, Funding acquisition

Frank Katzer: Conceptualization, Methodology, Formal Analysis, Resources, Writing-Review and Editing, Supervision, Project Administration, Funding acquisition

Mattie Christine Pawlowic: Conceptualization, Methodology, Formal Analysis, Resources, Writing-Original Draft, Supervision, Project Administration, Funding acquisition

## Acknowledgements

We would like to acknowledge the assistance of the Dundee Imaging Facility and University of Dundee Biological Services. Additionally, the Cellular Analysis and Molecular Analysis Shared Research Facilities at the College of Medical, Veterinary, and Life Sciences at the University of Glasgow. Additionally, the Bioservices department at Moredun Research Institute. We would like to acknowledge Lauren Lake, University of Dundee MSc Medical Art, for designing the calf illustrations included in Figures 1 and 2. We would like to thank Huvepharma for providing Parofor Crypto for the first two propagation studies conducted in calves. We would also like to thank Professors Susan Wyllie, David Horn, Marcus Lee and Dr. Megan Bergkessel for their advice and feedback during manuscript preparation.

This work was funded in part by a Sir Henry Dale Fellowship to MCP from the Wellcome Trust and the Royal Society (213469/Z/18/Z). This work was also funded in part by the Wellcome Trust via a Centre Award (203134/Z/16/Z and 223608/Z/21/Z), Innovator Award (223952/Z/21/Z), and an Institutional Strategic Support Fund (204816/Z/16/Z). This work was funded in part by the UKRI Biotechnology and Biological Sciences Research Council (BBSRC) grant, number BB/T00875X/1, via EASTBIO to Peyton Goddard and Sarah Stevens. Grant MJ Hall is supported by the Medical Research Council (MR/R015791/1). The staff and work conducted at Moredun were gratefully supported by the Scottish Government, Rural and Environment Science and Analytical Services Division (RESAS) through Underpinning National Capacity and the Moredun Foundation.

## Competing Interests

Huvepharma donated Parofor Crypto for propagation studies, namely passage 1 and 2 in neonatal calves. They sponsored FK to speak about *Cryptosporidium* at 2 events and a webinar organised by Huvepharma but neither the company nor their employees had any input in the study design, interpretation of the results for any studies, and were not involved in the preparation of this manuscript.

## Data accessibility statement

Raw data reported in complete in this manuscript. Materials will be made available to researchers as is possible, upon request to Dr. Mattie Christine Pawlowic.

