## Supplementary material for "Tissue atlas of *Cryptosporidium parvum* infection reveals contrasts between the natural neonatal calf model and laboratory mouse models": Goddard et al Supplemental

Goddard *et al.*, 2025.

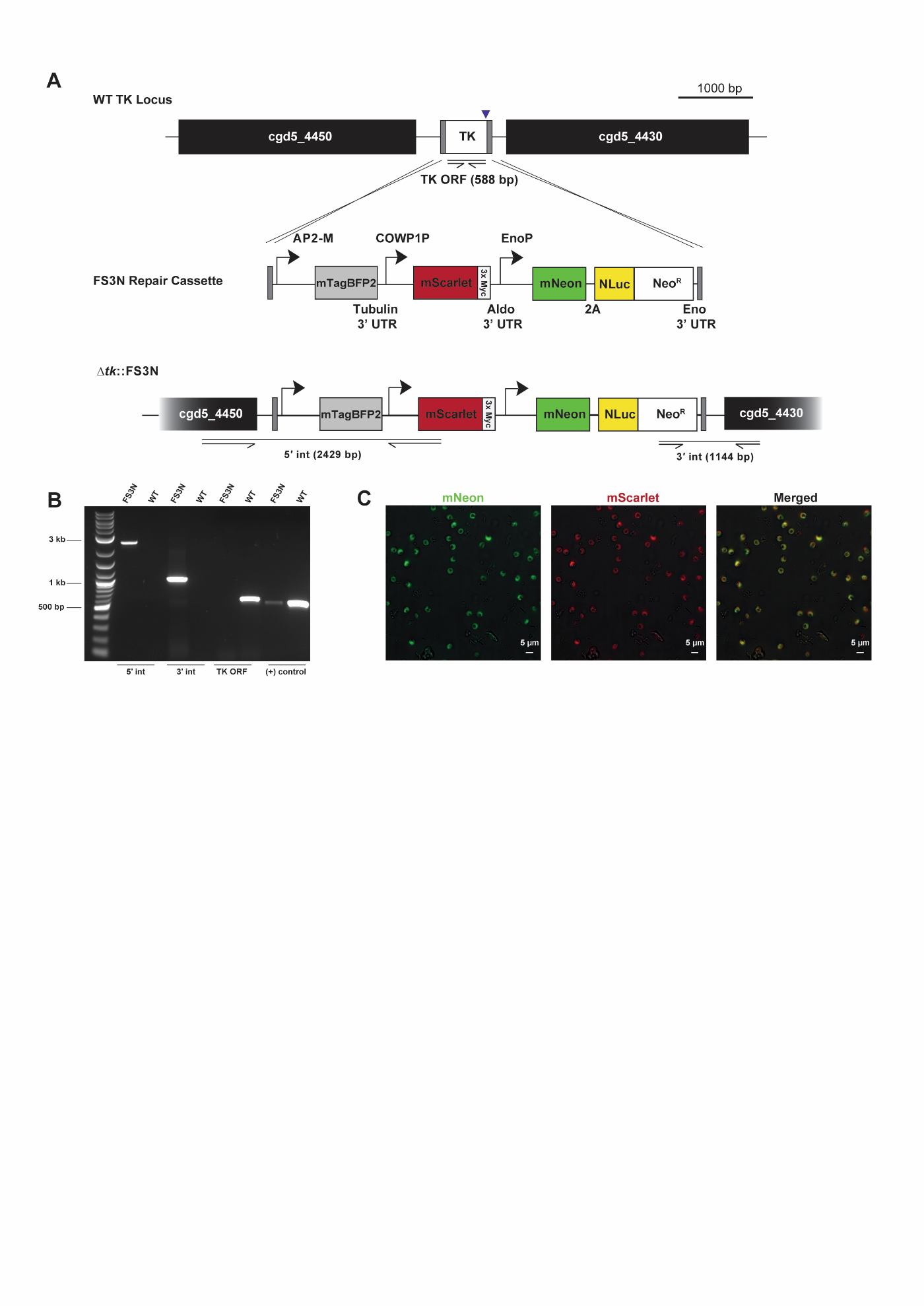

**Supplemental Figure 1. Generation & validation of transgenic *C. parvum* reporter strain.**

FS3N reporter parasites were designed to express fluorescent proteins during different parasite life cycle stages and were generated via the targeted deletion of the *C. parvum* thymidine kinase (TK) locus. (**A**) Schematic illustrating modification of the TK locus to produce the FS3N strain, *∆tk*::FS3N (*∆tk*::mScarlet-mNeon-Neo^R^). (**B**) Validation of correct integration of the FS3N construct by integration PCR. (**C**) Confocal imaging of live reporter oocysts, which express both mNeon (green) and mScarlet (red) fluorescent proteins in the cytoplasm.

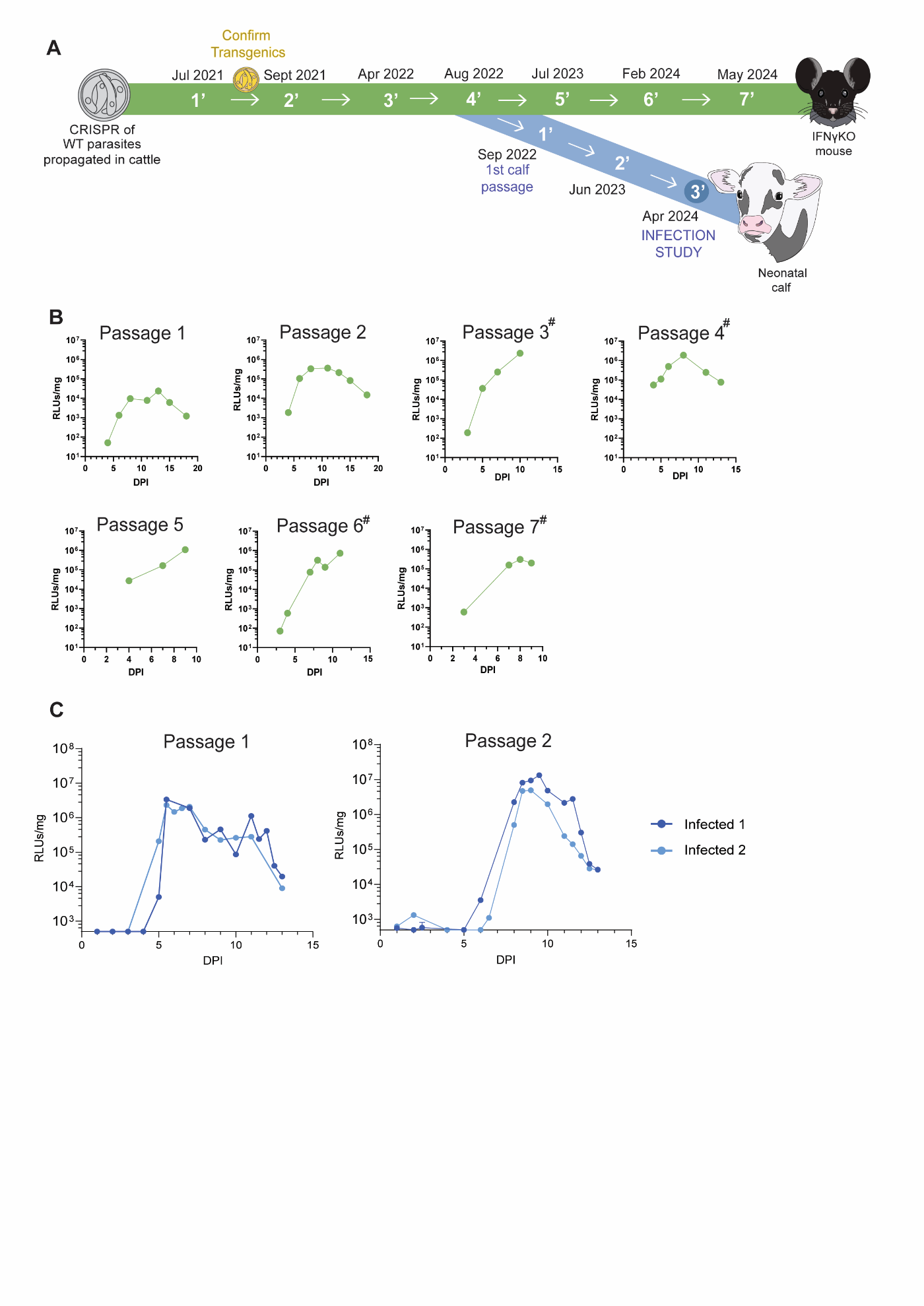

**Supplemental Figure 2. Propagation of reporter strain.**

Schematic indicating the timing and sequence of *in vivo* propagation of the reporter strain. (**A**) Reporter strain was generated in IFNγKO mice and then sequentially passaged in IFNγKO mice (blue) and neonatal calves (green). Parasites purified from the previous passage are used as the inoculum for the subsequent passage (**Supplementary Table 2**). Parasites purified from the fourth passage in IFNγKO mice (4’) were used as the inoculum for the first calf passage (1’). **B**) Faecal NLuc for each passage of reporter parasites in IFNγKO mice, plotted by cage. Faecal samples pooled from cage of 2 - 4 animals. RLUs/mg faeces ± SD; mean of 3 technical replicates plotted. ^#^Representative graph shown when more than one cage was infected in parallel. (**C**) Faecal NLuc for each passage of reporter parasites in neonatal calves, plotted by individuals. RLUs/mg faeces ± SD; mean of 3 or 6 technical replicates plotted. Where calves produced 2 samples a day, both are plotted; where 3 samples were produced the second and third were averaged for the second data point (Passage 2 Infected 2 DPI 8). Y-axis set at limit of detection (500 RLUs/mg). Data points less than or equal to the limit of detection were plotted as such.

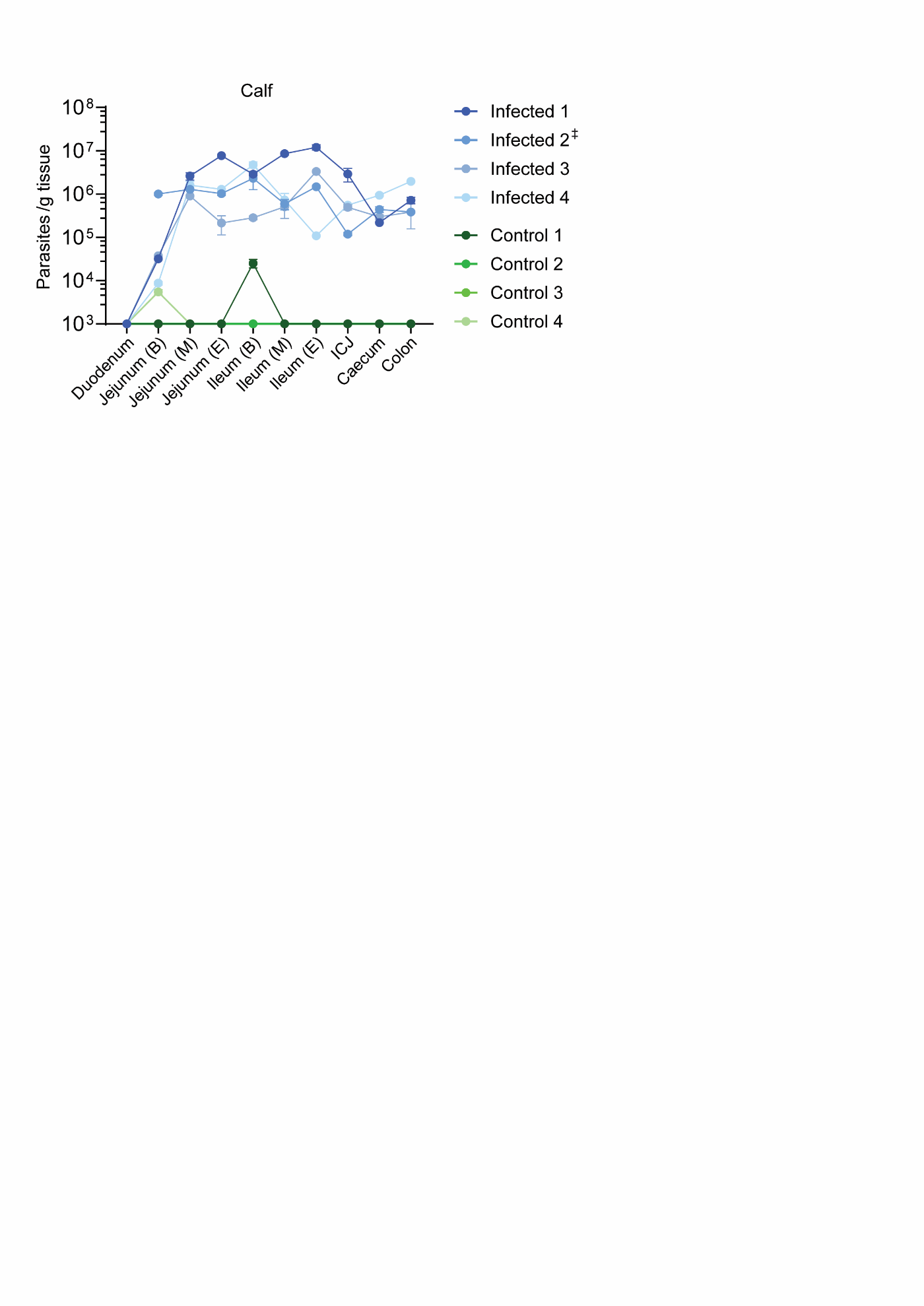

**Supplemental Figure 3. Infection of calf GI quantified by qPCR.**

Parasite burden in each tissue sample was quantified via qPCR. Parasites/g tissue ± SD mean of 3 technical replicates plotted. *Y*-axis begins at the limit of detection of 1x10^3^ parasites/g for clarity when plotting. All control calf samples were negative, with the exception of control calf 1 ileum and control calf 4 jejunum, which were lowly positive. ^‡^ “Infected 2” duodenum sample was not collected due to sampling error.

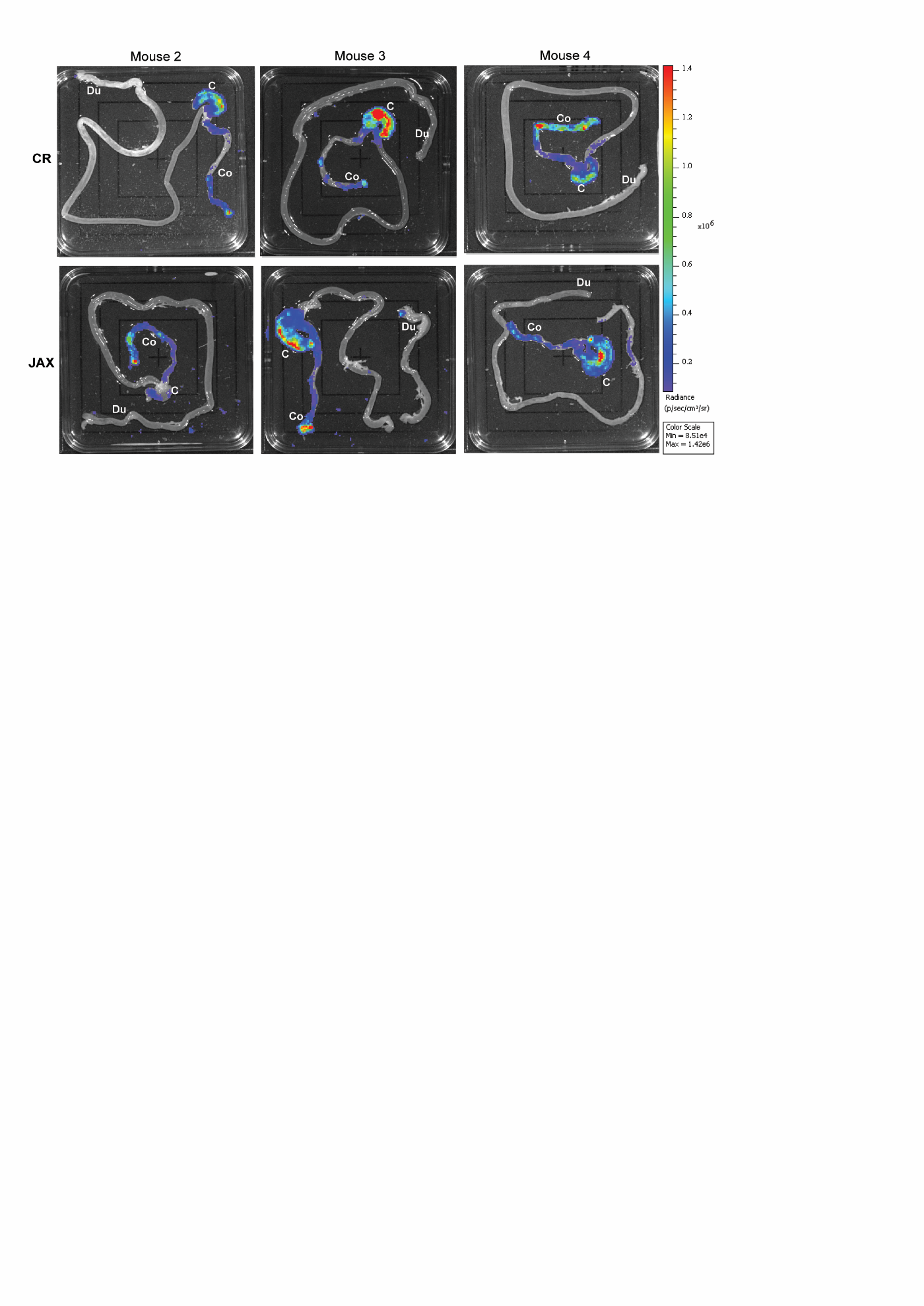

**Supplemental Figure 4. *Ex vivo* imaging of chronic NSG mice.**

*Ex vivo* imaging of GI from additional mice (representative image from mouse 1 in **Figure 3D** mice 2-4 here). Animals culled at peak infection. Animals purchased from different suppliers: “CR” Charles River, UK and “JAX” Jackson Labs, USA. Tissue locations of interest labelled: duodenum, Du; caecum, C; colon, Co.

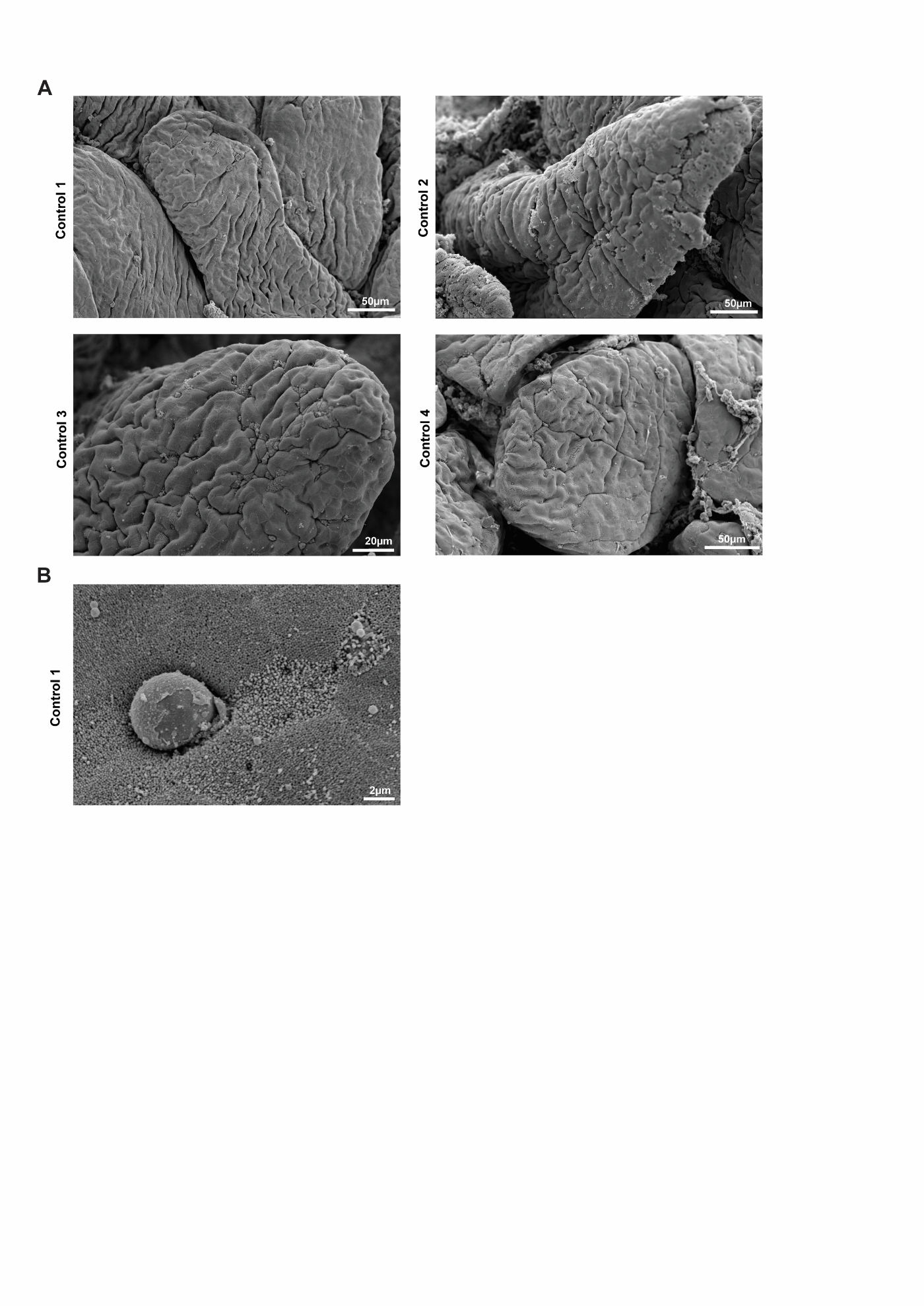

**Supplemental Figure 5. Scanning Electron Microscopy of control calf ileum.**

Tissue samples collected from the middle of the ileum of all control calves (Control 1 - 4) were imaged using scanning electron microscopy at low magnification. Normal villi morphology observed. **B**) A singular *Cryptosporidium* infection observed in Control 1, corresponding to detection of *C. parvum* by qPCR in the ileum of Control 1 (**Supplemental Figure 3**).

**Supplemental Table 1.** Oligonucleotides used in this study for generation and genetic validation of FS3N, tissue qPCR and GP60 genotyping.

| Purpose | Primer Name | Sequence (5' to 3’) |
| --- | --- | --- |
| F3SN Validation | F3SN 5’ Ins F | CTAATTCATCTGATTCTTCCCTC |
|  | F3SN 5’ Ins R | GTCTGCTAAGTATCTACCTCCGTCC |
|  | F3SN 3’ Ins F | GATCTTCTTTCTTCTCACCTTGCTCC |
|  | F3SN 3’ Ins | CAAAGTATCTTGATCTTTTGCTTCAC |
|  | TK ORF F | ATGGCAAAATTATACTTTTACTATTCAGCAATGAATGC |
|  | TK ORF R | TTAGAAATTGTATTCTTCACAATTAATTATATGATGTTTTCTGC |
|  | + control F  (α-tubulin) | CTAGTTATCCTTGTTCATTGAATTCTC |
|  | + control R  (α-tubulin) | TGAGCTCAAAAATATAAGATGGCAC |
| qPCR | *C. parvum* 18S-Foward | ATGACGGGTAACGGGGAAT |
|  | *C. parvum* 18S-Reverse | CCAATTACAAAACCAAAAAGTCC |
|  | *C. parvum* 18S-Probe | [FAM]-CGCGCCTGCTGCCTTCCTTAGATG-[BHQ1] |
| Genotyping | GP60 External F | ATAGTCTCCGCTGTATTC |
|  | GP60 External R | GAGATATATCTTGGTGCG |
|  | GP60 Internal F | TCCGCTGTATTCTCAGCC |
|  | GP60 Internal R | CGAACCACATTACAAATGAAG |

**Supplemental Table 2. Infection of transgenic *C. parvum* reporter strain in IFNyKO mice and neonatal calves.** Shedding pattern described in **Supplementary 2B and 2C** and **Figure 1B**.

| Group | Passage Number | Source of Inoculum | Date Infected | Age at time of Infection (Weeks) | Dose per animal | Excystation rate at time of infection (Calf samples only) |
| --- | --- | --- | --- | --- | --- | --- |
| Mouse | 1’ | WT | July 2021 | 6 | 1x10^7^ transfected sporozoites | - |
|  | 2’ | Mouse 1’ | Sept 2021 | 9 | 5x10^3^ oocysts | - |
|  | 3’ | Mouse 2’ | April 2022 | 11 | 5x10^3^ oocysts | - |
|  | 4’ | Mouse 3’ | August 2022 | 11 | 5x10^3^ oocysts | - |
|  | 5’ | Mouse 4’ | July 2023 | 39 | 1x10^4^ oocysts | - |
|  | 6’ | Mouse 5’ | Feb 2024 | 24 | 5x10^2^ oocysts | - |
|  | 7’ | Mouse 6’ | May 2024 | 15 | 1x10^3^ oocysts | - |
| Calf | 1’ | Mouse 4’ | Sept 2022 | Neonates | 5x10^6^ oocysts | - |
|  | 2’ | Calf 1’ | June 2023 | Neonates | 7.5x10^6^ oocysts | 50% |
|  | 3’ | Calf 2’ | April 2024 | Neonates | 7.5x10^6^ oocysts | 45% |

**Supplemental Table 3. Infection level of individuals, as indicated by faecal NLuc samples, at end of study.** Final NLuc activity measured from faecal sample of each animal prior to euthanasia (tissue NLucs reported in **Figure 2**). RLUs/mg is the mean of three technical replicates.

| Group | Animal | DPI | Mean NLuc (RLUs/mg) | Mean ± standard deviation |
| --- | --- | --- | --- | --- |
| Calf | Infected 1 | 8 | 3.96x 10^6^ | 1.91x 10^6^ ± 1.45 x10^6^ |
|  | Infected 2 | 8 | 1.48x 10^6^ |  |
|  | Infected 3 | 8 | 5.50x 10^5^ |  |
|  | Infected 4 | 8 | 1.63x 10^6^ |  |
| IFNγKO Mouse  (Tissue Atlas) | Infected 1 | 10 | 4.98x 10^5^ | 3.66x 10^5^ ± 2.83 x10^5^ |
|  | Infected 2 | 10 | 2.14 x 10^5^ |  |
|  | Infected 3 | 10 | 6.91x 10^5^ |  |
|  | Infected 4 | 10 | 5.91x 10^4^ |  |
| NSG Mouse  (Tissue Atlas) | Infected 1 | 11 | 8.26 x 10^5^ | 5.06x 10^5^ ± 2.78 x10^5^ |
|  | Infected 2 | 11 | 4.00 x 10^5^ |  |
|  | Infected 3 | 11 | 6.17 x 10^5^ |  |
|  | Infected 4 | 15 | 1.80 x 10^5^ |  |
| NSG Mouse  (*ex vivo* imaging) | CR *ex vivo* 1 | 34 | 2.82 x 10^6^ | 2.35x 10^6^ ± 9.35 x10^5^ |
|  | CR *ex vivo* 2 | 40 | 1.99 x 10^6^ |  |
|  | CR *ex vivo* 3 | 32 | 2.74 x 10^6^ |  |
|  | CR *ex vivo* 4 | 39 | 1.90 x 10^6^ |  |
|  | JAX *ex vivo* 1 | 39 | 3.87 x 10^6^ |  |
|  | JAX *ex vivo* 2 | 33 | 8.23 x 10^5^ |  |
|  | JAX *ex vivo* 3 | 40 | 1.71 x 10^6^ |  |
|  | JAX *ex vivo* 4 | 34 | 2.95 x 10^6^ |  |

**Supplemental Table 4. Calf feeding scores for tissue atlas study.** Feeding scores recorded twice a day: 0= animal feeding vigorously, 1 = complete feed with interruptions, 2 = complete feed with assistance, 3 = partial feed with assistance, 4 = refuses to feed. “- “indicates that no score was recorded. D8 no feed recorded as animals were euthanised AM.

| Group | Calf | Feeding Score | | | | | | | | | | | | | |
| --- | --- | --- | --- | --- | --- | --- | --- | --- | --- | --- | --- | --- | --- | --- | --- |
|  |  | DPI 1 | | DPI 2 | | DPI 3 | | DPI 4 | | DPI 5 | | DPI 6 | | DPI 7 | |
|  |  | AM | PM | AM | PM | AM | PM | AM | PM | AM | PM | AM | PM | AM | PM |
| Infected | 1 | 0 | 0 | 0 | 0 | 0 | 0 | 0 | 0 | 0 | 1 | 0 | 1 | 0 | - |
|  | 2 | 0 | 0 | 0 | 0 | 0 | 0 | 0 | 0 | 0 | 0 | 0 | 1 | 0 | 1 |
|  | 3 | 0 | 0 | 0 | 0 | 0 | 0 | 0 | 0 | 0 | 0 | 0 | 0 | 0 | 0 |
|  | 4 | 0 | 0 | 0 | 0 | 0 | 0 | 0 | 0 | 0 | 0 | 0 | 0 | 0 | 0 |
| Control | 1 | 0 | 0 | 0 | 0 | 0 | 0 | 0 | 0 | 0 | 0 | 0 | 0 | 0 | 0 |
|  | 2 | 0 | 0 | 0 | 0 | 0 | 0 | 0 | 0 | 0 | 0 | 0 | 0 | 0 | 0 |
|  | 3 | 0 | 0 | 0 | 0 | 0 | 0 | 0 | 0 | 0 | 0 | 0 | 0 | 0 | 0 |
|  | 4 | 0 | 0 | 0 | 0 | 0 | 0 | 0 | 0 | 0 | 0 | 0 | 0 | 0 | 0 |

**Supplemental Table 5. Calf demeanour scores.** Demeanour scores recorded twice a day: L = animal appears listless, R = reluctant to rise, U = unsteady on feet/ tottering gait, N = animal not remaining standing, “ / ” = animal healthy. “- “ indicates that no score was recorded. D8 no feed recorded as animals were euthanised AM.

| Group | Calf# | Demeanour Score | | | | | | | | | | | | | |
| --- | --- | --- | --- | --- | --- | --- | --- | --- | --- | --- | --- | --- | --- | --- | --- |
|  |  | Day 1 | | Day 2 | | Day 3 | | Day 4 | | Day 5 | | Day 6 | | Day 7 | |
|  |  | AM | PM | AM | PM | AM | PM | AM | PM | AM | PM | AM | PM | AM | PM |
| Infected | 1 | / | / | / | / | / | / | / | / | / | / | / | / | / | - |
|  | 2 | / | / | / | / | / | / | / | / | / | / | / | / | / | / |
|  | 3 | / | / | / | / | / | / | / | / | / | / | / | / | / | / |
|  | 4 | / | / | / | / | / | / | / | / | / | / | / | / | / | / |
| Control | 1 | / | / | / | / | / | / | / | / | / | / | / | / | / | / |
|  | 2 | / | / | / | / | / | / | / | / | / | / | / | / | / | / |
|  | 3 | / | / | / | / | / | / | / | / | / | / | / | / | / | / |
|  | 4 | / | / | / | / | / | / | / | / | / | / | / | / | / | / |

**Supplemental Table 6.** Mean tissue and faecal sample NLuc activity calculated from four independent experimentally challenged *C. parvum*-infected calves in the infection study (mean of the mean value from each of the 4 individual animals).

| Sample | NanoLuciferase activity | |
| --- | --- | --- |
|  | Mean | Standard Error |
| Duodenum | 818 | 281.9 |
| Jejunum (B) | 132,908 | 124,039.9 |
| Jejunum (M) | 1,217,750 | 443,132.7 |
| Jejunum (E) | 1,721,517 | 964,810.4 |
| Ileum (B) | 528,708 | 164,764.7 |
| Ileum (M) | 890,625 | 511,534.4 |
| Ileum (E) | 718,166 | 311,671.7 |
| Ileo-caecal Junction (ICJ) | 424,583 | 215,082.6 |
| Caecum | 109,459 | 17,083.5 |
| Colon | 368,442 | 215,849.4 |
| Faeces (D6) | 5,401,333 | 2,367,519.0 |
| Faeces (D8) | 3,810,834 | 1,452,633.0 |
